# Impacts of intraspecific variability and local adaptation on the ecophysiology of mosses: an example with *Sphagnum magellanicum*

**DOI:** 10.1101/859710

**Authors:** Tobi A. Oke, Merritt R. Turetsky, David J. Weston, A. Jonathan Shaw

**Affiliations:** Marine Science Institute, The University of Texas Austin, Port Aransas, TX, USA; Department of Integrative Biology, University of Guelph, Guelph, ON, Canada; Biosciences Division, Oak Ridge National Laboratory, TN, USA; Department of Biology, Duke University, Raleigh, NC, USA

**Keywords:** origin, local adaptation, phenotypic plasticity, intraspecific trait, morphological integration

## Abstract

**Background:** Bryophytes are a diverse plant group and are functionally different from vascular plants. Yet, plant ecology theories and hypotheses are often presented in an inclusive term. The trait-based approach to ecology is no exception; largely focusing on vascular plant traits and almost exclusively on interspecific traits. Currently, we lack information about the magnitude and the importance of intraspecific variability to the ecophysiology of bryophytes and how these might translate to local adaptation—a prerequisite for adaptive evolution.

**Method:** We used transplant and factorial experiments involving moisture and light to ask whether variability in traits between morphologically distinct individuals of *Sphagnum magellanicum* from habitat extremes was due to phenotypic plasticity or local adaptation and the implications for the ecophysiology of the species.

**Key Results:** We found that the factors that discriminated between the plant origins in the field did not translate to their ecophysiological functioning and the pattern of variability changed with the treatments, which suggests that the trait responses were due largely to phenotypic plasticity. The trait responses suggest that the need for mosses to grow in clumps where they maintain a uniform growth rate may have an overriding effect on responses to environmental heterogeneity, and therefore a constraint for local adaptation.

**Conclusion:** The circumstances under which local adaptation would be beneficial in this plant group is not clear. We conclude that extending the trait-based framework to mosses or making comparisons between mosses and vascular plants under any theoretical framework would only be meaningful to the extent that growth form and dispersal strategies are considered.

## Introduction

The importance of intraspecific trait variability to species persistence has long been appreciated by evolutionary biologists (Lande and Arnold, 1983; Schluter, 1988; Sides *et al*., 2014). Recent trait-based studies in ecology have also linked traits to community assembly, habitat preference and ecosystem functioning (Diaz *et al*., 2004; Cornwell *et al*., 2008; Pescador *et al*., 2015). Functional traits allow insights into plant-environment processes that are pertinent to community structure and function (Violle *et al*., 2007). However, unlike studies in evolutionary ecology, trait-based ecological studies more commonly focus on interspecific trait measurements, which implies that intraspecific variation in traits is less important to ecological processes (Bolnick *et al*., 2011). This focus on interspecific variation emphasizes the importance of species diversity on ecological processes but largely ignores how variation within populations affect plant performance and fitness. This importance of intraspecific trait variability has been discussed extensively (Bolnick *et al*., 2011; Violle *et al*., 2012; Siefert *et al*., 2015; Wright *et al*. 2016) but primarily from vascular plants perspective.

Bryophytes are a diverse plant group—comprising of 15000-20000 species (Shaw, Szövényi and Shaw, 2011) and are functionally different from vascular plants. Yet, plant ecological theories and hypotheses are often presented in an inclusive term. The trait-based ecology is no exception. Ecological inferences that may not necessarily apply to bryophytes are made, based on similarities in growth patterns and morphological and physiological configurations among vascular plants. Bryophytes lack complex morphological and physiological structures (e.g. roots and stomata) through which vascular plants actively interact with their environments for resource acquisition, conservation, and response to environmental heterogeneity (e.g. Hepworth *et al*., 2016). Even traits that seem comparable between bryophytes and vascular plants (e.g., leaf mass per area) are often difficult to quantify. However, bryophytes are capable of using facilitative interactions such as lateral movement (externally) of water across space (Rice, 2012) or vertical movement of water through their litter matrices, as a means of responding to change in their moisture environment. This cooperation for resource acquisition and retention means that individuals are buffered from the direct effect of the environment (e.g. Elumeeva *et al*., 2011). This highly integrated ecophysiological mechanism presents a rather unique inter- and intraspecific interactions where competition for resources such as moisture is weak (e.g. Hayward and Clymo 1982; Rydin 1985). Thus, intraspecific variability in this plant group may not necessarily have the same ecological meaning as it is understood for vascular plants.

Although there has been considerable research evaluating *Sphagnum* traits in the context of ecosystem function—particularly in linking *Sphagnum* species traits to aspects of water (Titus *et al*., 1983; Schipperges and Rydin, 1998; Hájek and Beckett, 2008) and carbon cycling (Turetsky *et al*., 2008; Laing *et al*., 2014; Bengtsson *et al*., 2016), only a few studies (Sastad and Flatberg, 1993; Sastad *et al*., 1999) have quantified intraspecific variability in traits. This is due to the difficulty in quantifying traits and perhaps also due to determining what constitutes an individual in clonal bryophytes like *Sphagnum*, because functional traits may only be measured at the level of an individual (Violle *et al*., 2007). However, viewing an individual as a structurally unattached, morphologically complete tissue—comprising of the capitulum, branch, and stem—the notion of individual is not complicated. That is, unattached individuals are physiologically independent and therefore, interact independently with their environments. Because moss tissues are not vascularized, they store a considerable amount of moisture externally (Elumeeva *et al*., 2011) and biomass likely influences the rate of moisture evaporation from their organs. Thus, examining the pattern of biomass investment into different organs and functions, such as the choice to invest in vertical growth versus branch mass, can be informative in evaluating the source, magnitude and the ecophysiological importance of variability in this plant group.

Variability often exists within a population because of sexual reproduction without apparent or immediate ecological benefits or consequences. Thus, intraspecific trait variability at the population level may reflect both the intrinsic genetic variability and phenotypic plasticity. One approach to evaluating the mechanistic importance of intraspecific variability is to explore trait variability in the context of local adaptation (Kawecki and Ebert, 2004). Plastic (non-genetic) responses to environmental heterogeneity could cause phenotypic differentiation within and among populations and this phenotypic differentiation may become genetically fixed by mutation and natural selection. Such differentiation on phenotypic responses to environmental heterogeneity is often the basis for local adaptation (Kawecki and Ebert, 2004). Additionally, locally adapted individuals would continue to exhibit adaptive responses that make them successful in their home environment even when they are subjected to a new environment where such response is no longer advantageous (Price *et al*., 2003; Kawecki and Ebert, 2004).

Investigating moss traits within the general framework of local adaptation can be informative in estimating the pattern, magnitude, and importance of intraspecific trait variability in this plant group.

Here, we explore the source, magnitude, and importance of intraspecific trait variability in *Sphagnum* moss. We ask whether there are differences in intraspecific trait variability and trait values between conspecifics from contrasting environments and whether the differences are due to adaptation to the conditions in their respective origin (hummock or hollow). That is, whether these differences are due to adaptive differentiation (local adaptation) or phenotypic plasticity. We focus on *S. magellanicum*, which is an ecologically dominant and widely distributed *Sphagnum* species. *S. magellanicum* is typically found in hollows and on low hummocks where moisture availability is high. However, it is also found within the carpets of *S. fuscum* on high hummocks—away from the water table, where a combination of high irradiation and moisture stress often impact photosynthesis and growth (Harley *et al*., 1989; Murray *et al*. 1993; McNeil and Waddington, 2003). The individuals of *S. magellanicum* found on hummocks often exhibit a reddish-brown pigmentation (as opposed to green), are less physically robust (e.g., slender stem and smaller capitulum) and relatively lower tissue water content compared with individuals found in hollows. This variation in phenotype is good for exploring intraspecific variability in the context of phenotypic plasticity versus local adaptation. Here, we capitalize on the pattern observed in the field to ask how intraspecific trait variability influence the breadth of environment where *S. magellanicum* is found. We test the following hypotheses.

- Since strong morphological integration (clump growth) of individuals is necessary for survival on hummock, which also promotes fast vertical growth, the hummock-originated individuals would consistently invest in vertical growth at the expense of biomass when grown in a common garden.
- Because green leaves tend to be more efficient for light capturing than red (anthocyanin-rich) leaves under low light (Burger and Edwards, 1996), we predict that hummock-originated plants would have lower photosynthetic potential under the shade treatment than hollow-originated plants. However, we expect the opposite when the plants are grown under full light on hummock because of the lack of pigmentation in the hollow-originated individuals.
- We hypothesize that hummock-originated individuals are locally adapted to low moisture availability and high irradiance that are prevalent in hummocks, we therefore predict that morphological and physiological responses of hummock-originated plants would be less sensitive to light and drought treatments compared with hollow-originated plants.

## Materials and Methods

In June 2016, we visited Wylde Lake bog in southern Ontario (43.91775, −80.40489) and collected individuals of *S. magellanicum* found on high hummocks, which are typically dominated by *S. fuscum* and thus represent an atypical environment for *S. magellanicum*. The sampling included plants from several hummocks because *S. magellanicum* is not typically found on hummocks and to collect enough samples for the experiments. Similarly, we collected individuals from hollow environments in which *S. magellanicum* was dominant. The *S. magellanicum* from hummocks are smaller and reddish-brown in colour whereas those from hollows were more physically robust and completely green. Hollow samples were kept separately from those collected from hummocks. All samples were immediately transferred to the University of Guelph phytotron where *S. magellanicum* samples from each environment were cut by knife into top 5 cm segments to exclude deeper, non-living component of the tissues and to create a standard length for all the plants.

### Hummock transplant experiment

We extracted four hummock monoliths, which comprised a continuous carpet of *S. fuscum* into surface peat to a depth of about 20 cm. The monoliths allowed us to incorporate the ecophysiological peculiarities (e.g. neighbourhood effect and vertical movement of moisture through litter matrices) of our study system into the experiment. Each monolith was gently placed in an 8.83-litre cylindrical pot. Each monolith was partitioned into equal halves with a stick, which was inserted horizontally into the surface of the moss carpet in each pot. Individuals of *S. magellanicum* from the two home environments (hummock versus hollow) were randomly assigned to a monolith and were inserted into the carpet of *S. fuscum*. Specifically, we inserted fifteen *S. magellanicum* hummock-originated individuals into one half of each monolith and fifteen hollow-originated individuals into the other half. Across the four replicate monoliths, we transplanted 60 plants from each plant origin. The hummock transplant experiment represents the breadth of “home” environment for individuals that were collected on hummocks in terms of substrate conditions, while hollow-originated plants in this case, were transplanted onto an “away” substrate. Two monoliths were assigned to a shade treatment and two were assigned to full light treatment. The shade treatment involved two shade boxes of 3.25 m x 1.47 m x 0.63 m in dimension and are built from PVC pipes. The shade boxes were covered with breathable 50% neutral density shade cloth. The 50% shade approximates the proportion of the photosynthetic active radiation (PAR) admitted into the *Sphagnum* carpet by the dominant vascular plant species (*Myrica gale*) at our site. This was obtained by measuring PAR below and above the canopy using the point sensor of a LI-250 light meter (LI-COR, Lincoln, Nebraska). These measurements were used to compute percentage of light admitted into the moss surface. The above canopy PAR ranged from 1206 – 2035 µmol m^-2^ s^-1^ whereas below canopy values ranged from 224 −1714 µmol m^-2^ s^-1^. We did not find a difference in the moisture profiles of hummocks sampled along moisture gradient in our site, therefore, we did not vary moisture for this experiment.

### Factorial light x moisture experiment

Our second experiment involved a 3 × 2 factorial experiment with two plant origins (hummock versus hollow), two light treatments (full light; 50% light) and two water treatments (saturated; low water). This experiment represents the breadth of “home” environment for hollow-originated individuals in terms of substrate conditions, while hummock-originated plants in this case were transplanted onto “away” substrates. The shade treatment was imposed as described above. The drought treatment was created by maintaining treatment pots at an average volumetric water content of about 12%, which is the mean summer volumetric water content at the top 1 cm of moss in the field site. The saturated water treatment was maintained by monitoring and topping up the experimental pots with water, and volumetric water content consistently exceeded 21%. The water contents across all experimental pots were monitored with a portable Hydrosense soil moisture meter (Campbell Scientific, Inc., USA).

The experimental pots were filled with 3 cm of deep peat moss underneath a 1 cm layer of surface peat. The deep peat was from a commercial source while the surface peat was extracted from the field in an area near the *Sphagnum* collections in hollow. The pots were 227.4 cm^3^ in size, with holes at the base through which water was fed into the pots. There were 9 plants (one plant per pot) for each of the four treatment combinations (9 plants x 2 origin x 2 light x 2 moisture treatments), which we replicated twice. Thus, a total of 144 plants were used in the experiment. Because bogs are nutrient-poor and typically fed by rainwater, the plants were not fertilized and were watered exclusively with rainwater that was harvested in Guelph.

In the context of local adaptation, each experiment contains aspects of a “home” versus an “away” treatment (Kawecki and Ebert, 2004; Blanquart *et al*., 2013). In the first transplant experiment, hummock individuals transplanted onto the hummock mesocosms represent a “home” treatment while hollow individuals represent an “away” treatment. However, this transplant experiment is an incomplete design but it was not possible for us to maintain hollow mesocosms due to the extremely unconsolidated (low bulk density) nature of hollow surface soils and species homogeneity. The combination of the experiments nonetheless represents a range of environment that the species is typically exposed to and allows us to at least reduce the confounding effects of unmeasured environments (Kawecki and Ebert, 2004).

### Quantification of traits

The two experiments ran fully from July 2016 to January 2017. At the end of the experiments, we measured a suite of traits on individuals from each treatment. We quantified two traits related to growth, including vertical growth rate (growth per time) and biomass. We also measured allocation of biomass into capitulum, branch, and stem. The capitulum is taken as the top 1 cm of the plant (Clymo 1970). Branch mass was determined by removing, drying and weighing the stem, leaves and branches (fascicles), which were collectively measured as branch mass. The exposed stem after removal of capitula and branches was dried and weighed to obtain stem mass.

We also quantified the dark respiration as a measure of metabolic activity. Respiration rates were measured on six individuals per treatment, which were selected at the end of the experiment. For these individuals, we placed the entire plant in a dark glass jar. The jars were sealed with stopcocks and placed under their respective treatment environment. The CO_2_ in the jar headspace was drawn three times at 3 hr intervals with gas-tight syringes. The CO_2_ concentration was analyzed with an EGM-4 infrared gas analyzer (PP Systems, Hitchin, Hertfordshire, UK). We performed linear regressions of CO_2_ concentration against time, using the slopes of these relationships as our measurement of respiration rate. We then used the dry mass of the samples to convert the slopes into µmol of CO_2_g^-1^minute^-1^.

Finally, we measured the dark-adapted fluorescence (F_v_/F_m_) as a measure of maximal photosynthetic efficiency. The dark-adapted F_v_/F_m_ measurements were taken at the end of the experiment. Individuals from each treatment were placed in the dark for at least 6 hours to ensure that Q_A_ electron acceptors are fully reduced and that reaction centers are in the ‘open’ state. We then quantified dark-adapted F_v_/F_m_ on each plant using a pulse-modulated fluorometer (OS1p, Opti-Sciences, Hudson, NH).

## Statistical analyses

The data were explored for normality and where there was a departure from normality (vertical growth rate and branch mass), they were transformed using a logarithm transformation. Because the plants in the hummock transplant experiment were grown in only four pots, we tested for differences in trait values using mixed effect models, where we analyzed pot ID as a random effect to account for lack of independence. Multiple mean comparisons were obtained for models with interaction effects using “lsmeans” package in R. We tested for mean trait values in the factorial experiment using 3-way ANOVA and obtained multiple mean comparisons for interaction effects using Tukey HSD. We explored patterns of trait variability across experimental treatments by partitioning the variance in the data using the varpart function in R package “Vegan”. We used this approach combined with redundancy analysis to examine how the experimental treatments influenced within-trait variability and total trait variability. All analyses were performed in R 3.2 (R core Development Team 2015) and all statistical tests were conducted at α = 0.05.

## Results

### Hummock-transplant experiment

Hummock-originated plants had lower F_v_/F_m_ than hollow plants (Fig. 1a) with no other significant main effects or interactions (Table 1). Vertical growth rate, capitulum mass, and respiration were consistently higher under the shade than the high light treatment (Fig. 1b & c). Total biomass and stem biomass was influenced by a plant origin x light interaction (Fig 1d). Hummock plants tended to have lower total and stem biomass than hollow plants but only in the shade treatment.

**Table 1.** Results of a mixed effect model for the hummock transplant experiment showing F and p-values specific to each trait. Bold texts are significant values (p < 0.05). O = Origin, L = Light and DF = treatment and sample degrees of freedom

| Treatments | DF | Respiration |  | Vertical | Total | Capitulum | Branch | Stem |
| --- | --- | --- | --- | --- | --- | --- | --- | --- |
| | | ($\mu\text{mol}^{-1}\text{g}^{-1}\text{min}$) | $F_v/F_m$ | growth rate<br>(cm) | Biomass<br>(g) | mass<br>(g) | mass<br>(g) | mass<br>(g) |
| O | 1, 59 | 0.21, 0.651 | **4.4, 0.040** | 2.4, 0.128 | 2.6, 0.115 | 1.3, 0.267 | 0, 0.915 | **6.8, 0.012** |
| L | 1, 59 | **9.2, 0.038** | 1.3, 0.264 | **50.2, <0.0001** | **12.4, 0.026** | **21.1, <0.0001** | 0.77, 0.382 | 3.1, 0.146 |
| O*L | 1, 59 | 1.5, 0.222 | 0.1, 0.764 | 2.5, 0.122 | **7.1, 0.010** | 3.9, 0.053 | 1.9, 0.172 | **4.1, 0.048** |

**Fig. 1.**
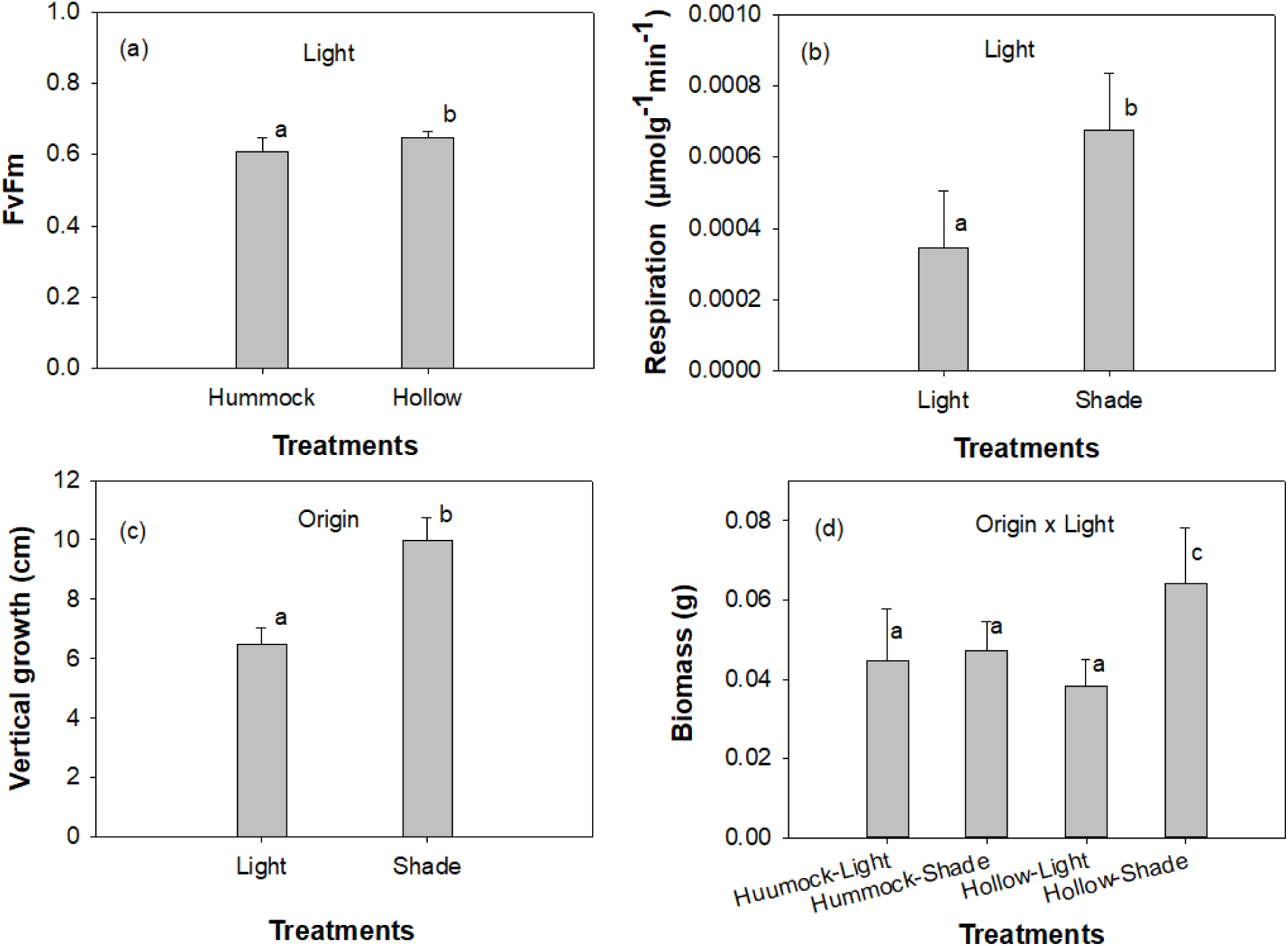
Results of mixed effects models examining trait variation to treatments in hummock transplant experiments (a) F_v_/F_m_ averaged by plant origin (b) respiration averaged by light treatment (c) vertical growth averaged by light treatment (d) biomass averaged by a light x plant origin treatment interaction. Same letter notation depicts no differences between means based on Tukey HSD post-hoc tests.

We found strong positive correlations between some of the traits. There were correlations for example between vertical growth rate and respiration rate and between respiration rate and biomass for both hummock and hollow plants (r^2^ = 0.24, p < 0.05 and r^2^ = 0.56, p < 0.001) and hollow plants (r^2^= 0.30, p < 0.05 and r^2^ = 0.73, p < 0.001) (Fig. 2a & b). However, the effect of plant origin on these relationships was not statistically significant (p > 0.05).

**Fig. 2.**
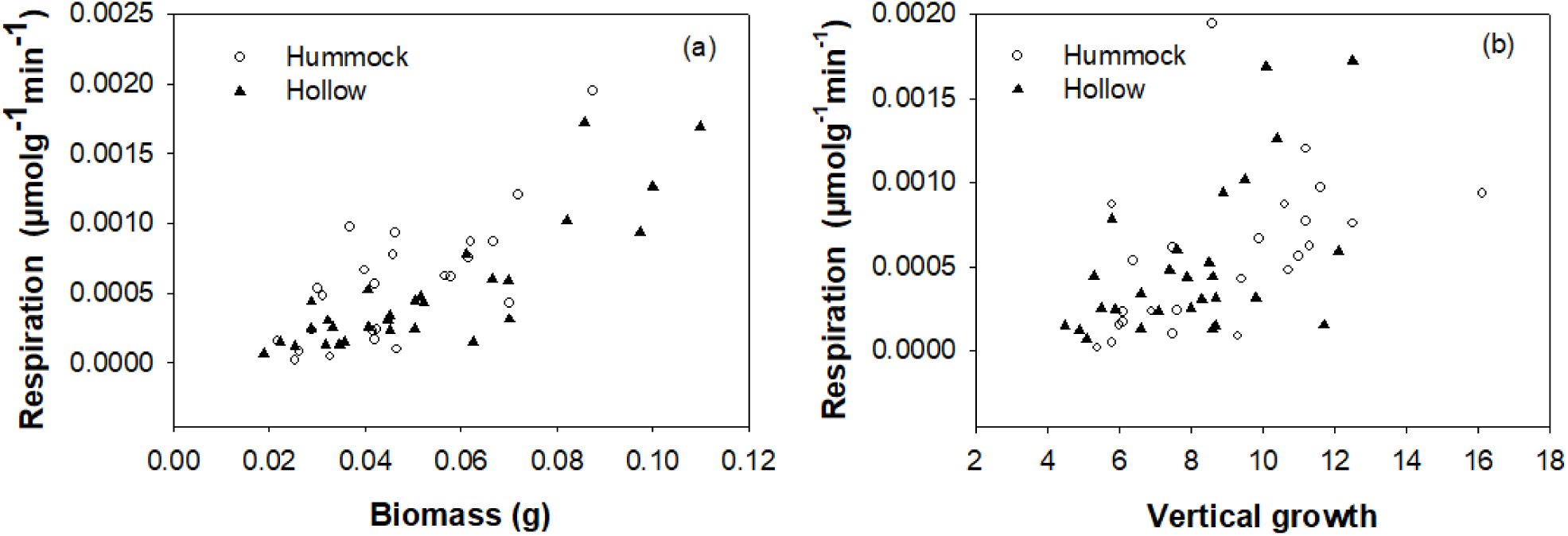
Correlational relationships between respiration and total biomass (a) and the relationship between respiration and vertical growth rate (b) in the hummock transplant experiment.

For most traits, plant origin did not explain a significant amount of variation in individual traits (0 −10%), while light explained between 0 - 46% (Table 2). Origin (hummock vs. hollow) explained significant variation for F_v_/F_m_ and stem mass, while light explained significant variation in vertical growth, capitulum mass, and total biomass (Table 2). When analyzed for total variability across all traits (respiration, F_v_/F_m_, capitulum, branch, stem, and total biomass), plant origin only accounted for 2% of the variability (p > 0.05) whereas light accounted for 16% (p < 0.001).

**Table 2.**
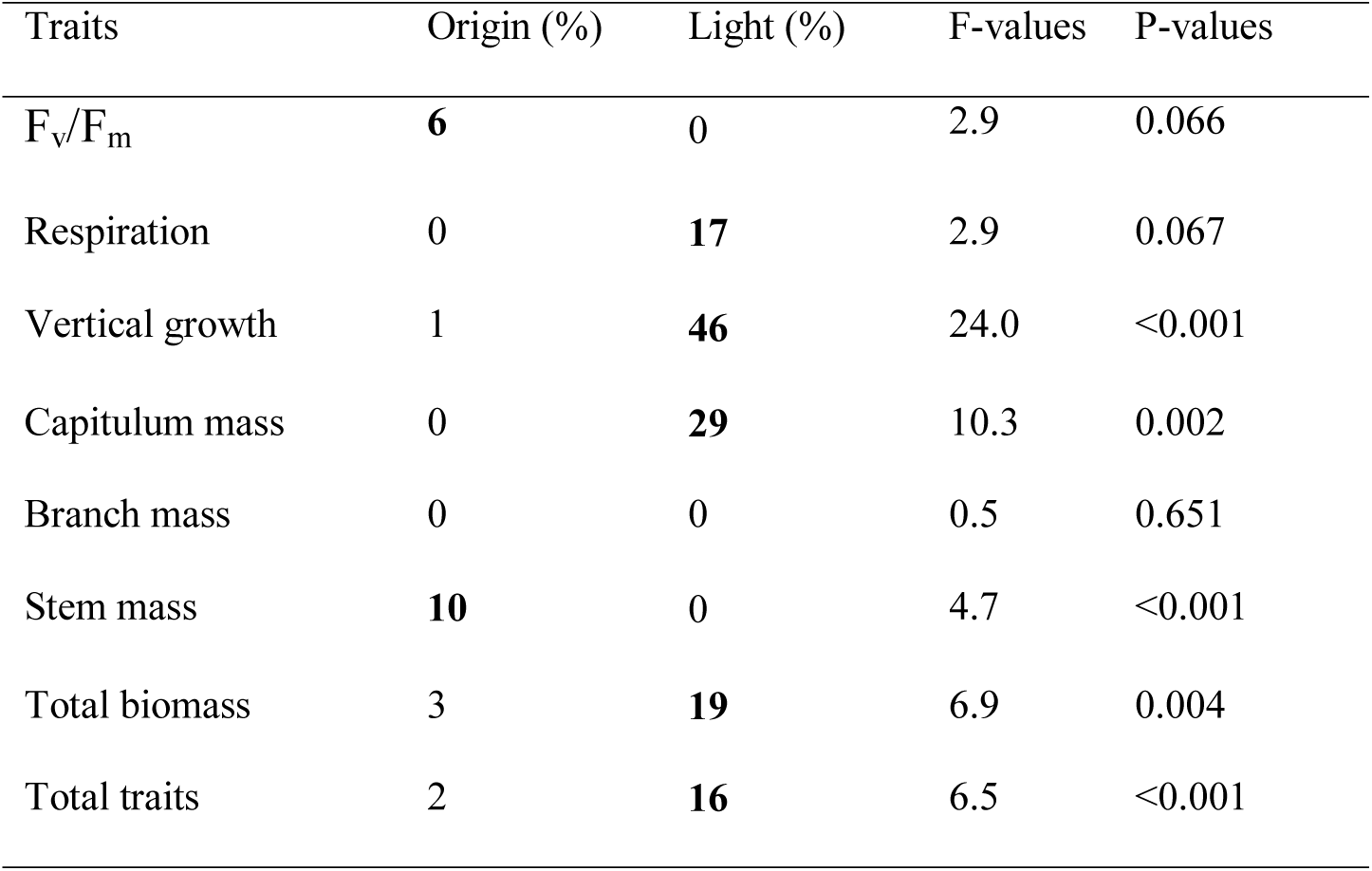
The effects of plant origin and light treatment on individual trait variability as well as total trait variability for hummock transplant experiment. Bold figures are statistically significant trait variability values (p < 0.05) under each parameter while the p-values are the overall p-values of the model.

### Light x moisture factorial experiment

In the factorial experiment, traits were more generally influenced by the main effects of origin and moisture than their interaction effects or the main effect of light (Table 3). The post-hoc tests showed that capitulum mass was greater in hummock plants than in hollow plants under the high moisture treatment (p < 0.05) but did not significantly differ between the plant origins under the low moisture treatment. The opposite trend was true for branch mass as hollow plants had a greater branch mass than hummock plants under the high moisture treatment (p < 0.05) but there was no difference in branch mass between the origins under the low moisture treatment. Interestingly, the stem mass of hollow plants subjected to low moisture was greater than stem mass of hummock plants subjected to high moisture (p < 0.001). Vertical growth rate was fastest under the high moisture treatments regardless of light (Fig. 3a) and was lower under the low moisture treatments. Biomass was greatest at the high light and high moisture treatment and tended to be lowest under the low moisture treatments across both light treatments (Fig. 3b). F_v_/F_m_ was higher in hollow individuals than in hummock individuals, especially under high moisture (Fig. 3c). Respiration was higher under high moisture than the low moisture treatment and did not vary with light (Fig. 3d).

**Table 3.**
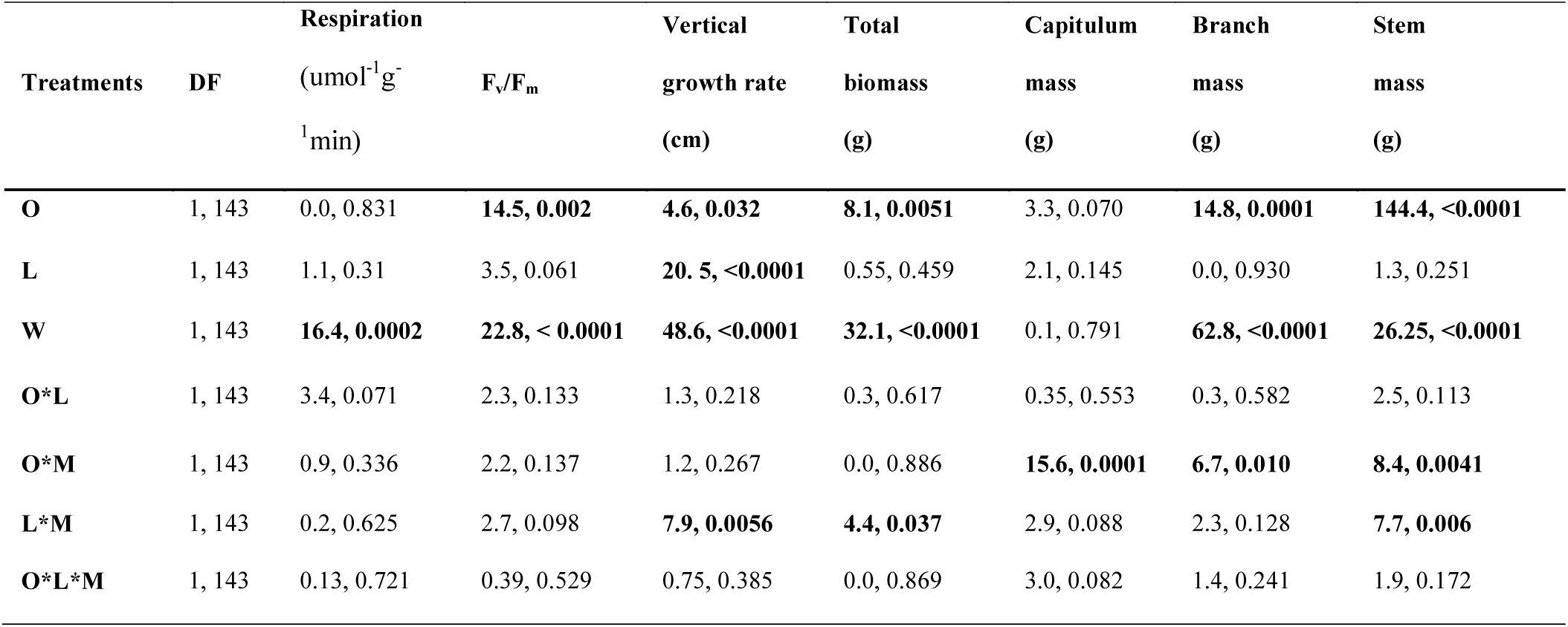
Results of a 3-way ANOVA for the factorial experiment showing F and p-values for the traits. Bold texts are significant values (p < 0.05). O = Origin, L = Light, M = Moisture and DF = treatment and sample degrees of freedom

| Treatments | DF | Respiration |  | Vertical | Total | Capitulum | Branch | Stem |
| --- | --- | --- | --- | --- | --- | --- | --- | --- |
| | | ($\mu\text{mol}^{-1}\text{g}^{-1}\text{min}$) | $F_v/F_m$ | growth rate<br>(cm) | biomass<br>(g) | mass<br>(g) | mass<br>(g) | mass<br>(g) |
| **O** | 1, 143 | 0.0, 0.831 | **14.5, 0.002** | **4.6, 0.032** | **8.1, 0.0051** | 3.3, 0.070 | **14.8, 0.0001** | **144.4, <0.0001** |
| **L** | 1, 143 | 1.1, 0.31 | 3.5, 0.061 | **20.5, <0.0001** | 0.55, 0.459 | 2.1, 0.145 | 0.0, 0.930 | 1.3, 0.251 |
| **W** | 1, 143 | **16.4, 0.0002** | **22.8, <0.0001** | **48.6, <0.0001** | **32.1, <0.0001** | 0.1, 0.791 | **62.8, <0.0001** | **26.25, <0.0001** |
| **O\*L** | 1, 143 | 3.4, 0.071 | 2.3, 0.133 | 1.3, 0.218 | 0.3, 0.617 | 0.35, 0.553 | 0.3, 0.582 | 2.5, 0.113 |
| **O\*M** | 1, 143 | 0.9, 0.336 | 2.2, 0.137 | 1.2, 0.267 | 0.0, 0.886 | **15.6, 0.0001** | **6.7, 0.010** | **8.4, 0.0041** |
| **L\*M** | 1, 143 | 0.2, 0.625 | 2.7, 0.098 | **7.9, 0.0056** | **4.4, 0.037** | 2.9, 0.088 | 2.3, 0.128 | **7.7, 0.006** |
| **O\*L\*M** | 1, 143 | 0.13, 0.721 | 0.39, 0.529 | 0.75, 0.385 | 0.0, 0.869 | 3.0, 0.082 | 1.4, 0.241 | 1.9, 0.172 |

**Fig. 3.**
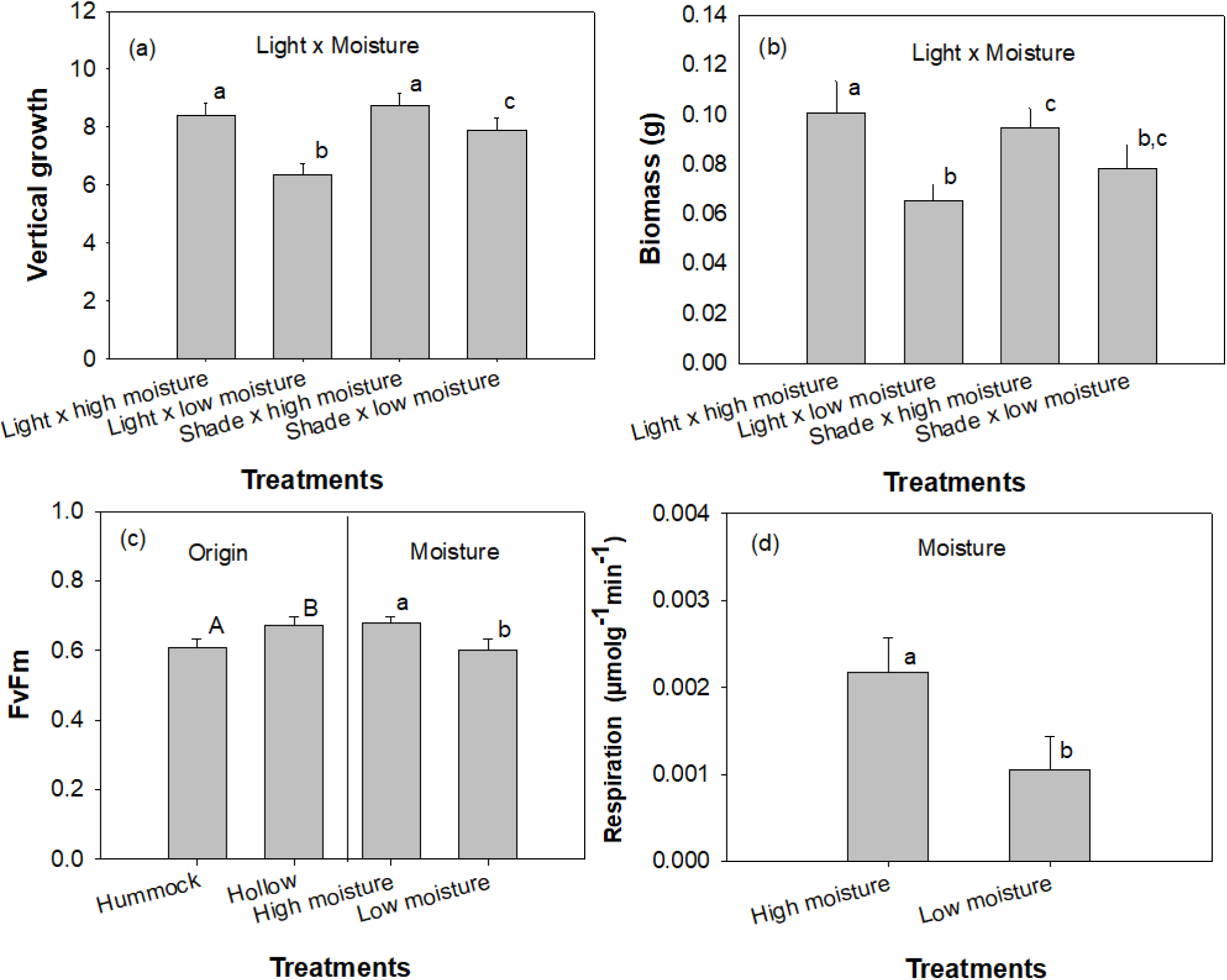
Effects of treatments in the factorial experiment on *S. magellanicum* traits (a) vertical growth averaged by a light x moisture treatment interaction, (b) biomass averaged by a light x moisture treatment interaction, (c) canopy F_v_/F_m_ averaged by plant origin and moisture treatments, and (d) respiration averaged between the moisture treatments. Same letter notation depicts no differences between means based on post hoc tests. There was no origin x light interaction on F_v_/F_m_.

Consistent with the hummock transplant experiment, we found strong positive correlations between respiration and biomass and between respiration and vertical growth for both hummock (r^2^ = 0.25, p < 0.001 and r^2^ = 0.57, p < 0.001) and hollow (r^2^= 0.53, p < 0.001 and r^2^ = 0.65, p < 0.001) plants (Fig. 4a & b). However, unlike in the hummock transplant experiment, the pair-wise comparison of the relationships between respiration and biomass was influenced by plant origin (z = 2.65, p = 0.01).

**Fig. 4.**
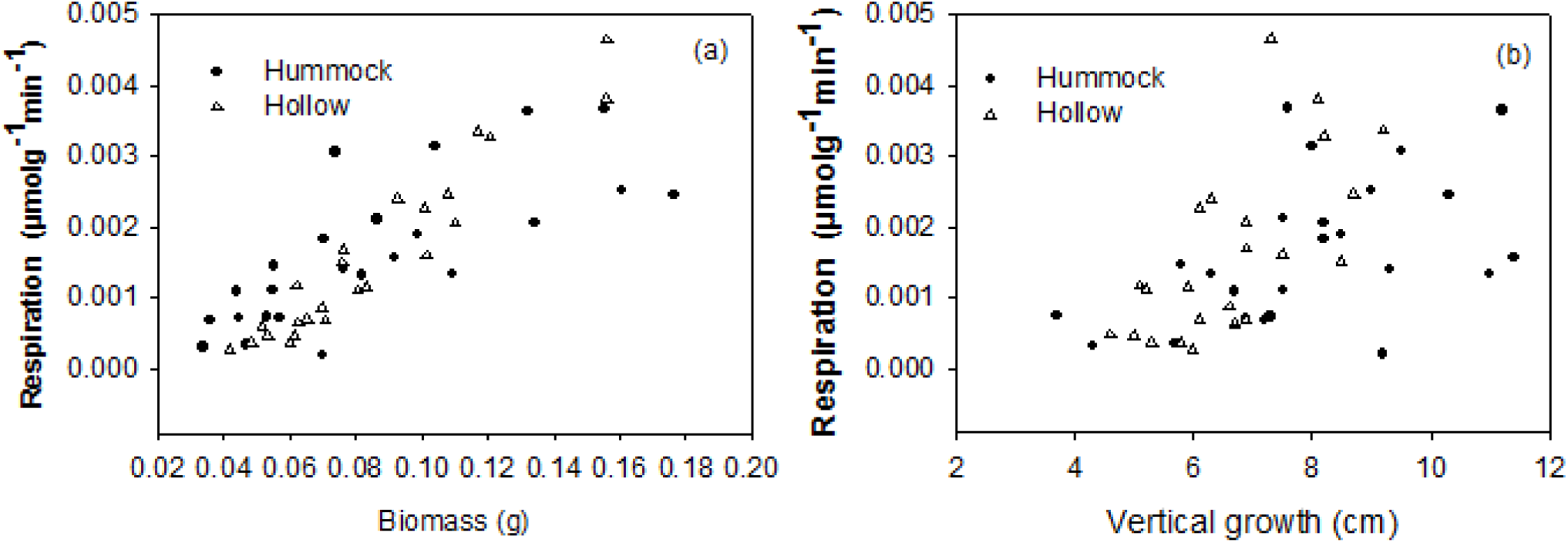
Correlational relationships between respiration, biomass and vertical growth in the factorial experiment for hummock and hollow originated plants.

Plant origin explained the most variation in stem mass (44%) relative to moisture and light. In general, the influence of the light treatments explained little or no variation among traits in this experiment. Moisture explained a significant amount of variation in all traits except for capitulum mass and was particularly important for respiration and branch mass variation. We found that plant origin and moisture explained similar levels of total variation across traits (Table 4). The data were also split into two independent datasets based on plant origin and were accordingly explored for variability due to light and moisture effects. Light explained 1% of total variability in hollow plant traits and 4% in hummock plant traits whereas moisture explained 22% of variability in hollow plant traits and 13% in hummock plant traits. However, the effect of light on variability of hollow plant traits was not statistically significant.

**Table 4.** Percentage trait variability due to plant origin as well as experimental light and moisture treatments in the factorial experiment. Bold figures are statistically significant trait variability values (p < 0.05) under each parameter while the p-values are the overall p-values of the model.

| Traits | Origin (%) | Light (%) | Moisture (%) | F-values | P-values |
| --- | --- | --- | --- | --- | --- |
| $F_v/F_m$ | **7** | 1 | **12** | 12.1 | <0.001 |
| Respiration | 0 | 0 | **26** | 5.7 | 0.005 |
| Vertical growth | 2 | **8** | **21** | 22.0 | <0.0001 |
| Capitulum mass | 2 | 0 | 0 | 2.0 | 0.112 |
| Branch mass | **6** | 0 | **26** | 21.3 | <0.001 |
| Stem mass | **44** | 0 | **7** | 46.6 | <0.001 |
| Total biomass | **3** | 0 | **15** | 10.8 | <0.001 |
| Total traits | **11** | 1 | **14** | 16.5 | <0.001 |

## Discussion

Understanding the environmental mechanisms influencing trait variability is important for assessing not only how the environment influences species distribution but also how species influence ecosystem function (de Bello *et al*., 2011; Mitchell and Bakker, 2014; Bennett *et al*., 2016). Our goal was to evaluate the magnitude of variability within and among traits, and the response of traits to varying environmental mechanisms. This is not an attempt to characterize *Sphagnum* physiology but rather to explore the importance of trait variability in controlling responses to environmental heterogeneity.

### Effect of plant origin (hummock versus hollow) on Sphagnum traits

Although many bryophytes are clonal, studies have shown that their populations can be as genetically diverse as any population of non-clonal vascular plants (Stenoien and Sastad, 1999) and *Sphagnum* does show spatial patterns of genetic variability (Gunnarsson *et al*., 2007). Given our sampling design, we assumed that our hollow and hummock-originated plants represent genetically disparate groups and that the differences in their phenotypic traits are indicative of adaptive differentiation (local adaptation). The generally weak effect of plant origin on the traits in the hummock transplant experiment relative to that in the factorial experiment suggests that the trait responses were largely due to phenotypic plasticity as opposed to local adaptation.

A unique characteristic of *Sphagnum* is that it acquires and conserves moisture through stem and canopy integration (clump growth form), especially on hummocks. That is, individuals may not grow considerably taller than their neighbors without experiencing desiccation (Hayward and Clymo, 1983). Thus, *S. magellanicum* growing on hummocks may not grow considerably faster or taller than the typical height of *S. fuscum*-derived carpet. Pure stands of *S. magellanicum* typically grow faster than those of *S. fuscum* (Breeuwer *et al*., 2008), which implies that *S. magellanicum* plants (regardless of their origin) growing on *S. fuscum*-dominated hummocks may not express their maximum growth rates. This likely explains why plant origin did not influence traits in the hummock transplant experiment.

Environmental heterogeneity may cause phenotypic differentiation that may not translate into genetic differentiation (Kawecki and Ebert, 2004). Such non-genetic phenotypic heterogeneity likely explains the phenotypic differences that were observed between the plant origins in the field. The major differences between our experiments and between the plant origins in the field are the distance to water table and interspecific clumping (morphological integration) in the hummock transplant experiment. It is not clear however, whether the effects of origin in the factorial experiment are indicative of phenotypic differentiation that was observed between the plant origins in the field. Because if morphological integration indeed overrode the expression of phenotypic differentiation on hummock, then the clump growth form of *Sphagnum* would likely constrain local adaptation.

### Light controls on Sphagnum trait variation

Light (through UV damage) is a common stressor influencing bryophytes’ performance (Post *et al*., 1990; Marschall and Proctor, 2004). Although many *Sphagnum* species including *S. magellanicum* grow in open habitats, photoinhibition is known to constrain photosynthetic processes of *Sphagnum* species (Murray, Tenhunen and Nowak, 1993; Hájek, 2014) and it is perhaps more common in hummock habitats where low canopy moisture and high irradiance are prevalent (Bragazza, 2008). Since typical hummock species are rarely completely green (unless under shade), we hypothesized that the reddish pigmentation found in hummock plants is for photoprotection (Bonnett *et al*., 2010) and therefore is an adaptive trait for colonizing or surviving in hummocks. This is plausible if photoprotection prevents photoinhibition and therefore, would lead to higher photosynthetic potential in hummock plants compared with hollow plants under full light. This could lead to low light capture and low photosynthetic potential when growing under shade as has been observed in vascular plants (Burger and Edwards, 1996). Contrary to our predictions, our results showed that the hummock plants had relatively lower F_v_/F_m_ across all experimental treatments compared with the hollow plants. Also, under the shade treatments, some of the hummock plants changed from reddish to light pink colour and some with a tint of green, which is consistent with the findings that pigmentation of *S. magellanicum* is plastic (Yousefi *et al*., 2017). This means that the pigmentation is induced by the environment. The generally low F_v_/F_m_ in hummock plants suggests that the reddish pigmentation might prevent an optimal photosynthetic response, which would mean that there is a cost to achieving photoprotection. However, we did not find any relationship between F_v_/F_m_ and total biomass, which is often used as a proxy for fitness in plants (Younginger *et al*., 2017).

Shade tends to reduce transpiration (Muthuchelian *et al*.,1989; Pons *et al*. 2001; Gent, 2007), which would diminish the need for morphological integration. Under the shade treatment of the hummock transplant experiment, the plants were more robust (e.g., bigger capitulum) and the moss canopy was generally rough and loose compared with light treatment, which was relatively smooth and compacted. This disparity in growth response due to the difference in light level likely contributed to the strong effect of light on trait variability in the hummock transplant experiment. Surprisingly, light was less important to trait variation in the factorial experiment. This could be because we only manipulated moisture in the factorial experiment, which is well established as having an important role in *Sphagnum* growth and distribution (McNeil and Waddington, 2003; Oke and Hager, 2017) and also was the dominant source of trait variation in the factorial experiment.

### Implications of trait variability and local adaptation in Sphagnum

Trait variability is considered one of the mechanisms by which plant populations cope with environmental heterogeneity (Jung *et al*., 2014) and it is deemed the raw material for natural selection (Bolnick *et al*., 2011). For instance, high trait variability could help a population to transition to a new trait optima and therefore to a new adaptive peak through natural selection (Bürger, 1999). In this study, most of the variability remained unexplained by our treatments. However, it is important to note that most traits measured in this study exhibited low levels of variation. It is also important to note that clonality is common in *Sphagnum*, especially at fine scales, which may lead to low phenotypic variation. Low phenotypic variation may be advantageous for morphological integration. Although our sampling design was intended to avoid repeatedly sampling clones, it is not uncommon for a *Sphagnum* population to be dominated by a single clone (Cronberg, Molau and Sonesson, 1997; Gunnarsson, Shaw and Lonn, 2007), which would then likely be overrepresented in our experiments.

Due to the generally low nutrient condition that limits spore germination in peatlands (Sundberg & Ryadin 2008), *Sphagnum* populations are maintained largely by clonal growth (Cronberg *et al*., 1997; Gunnarsson *et al*., 2007). That is, dispersal by spore in *Sphagnum* is long-distant and random (Whitaker and Edwards, 2010). This is true for many moss species (Miles and Longton, 1992), which means that there is a low accruable benefit in passing down the local selective advantage through spores. While the short-distance dispersal through clonal growth is less random, it likely perpetuates homogeneity of trait. Trait homogeneity may have an ecophysiological value in stem and canopy integration for moisture retention and survival. However, as observed in the field and as demonstrated in this study, morphological integration is quite common in *Sphagnum* even among species with different growth rates. This means that stem and canopy integration is more likely a function of plasticity rather than trait homogeneity. Thus, given their mode of dispersal and the clump growth form, locally adapted growth responses may not be beneficial to mosses. In any case, extending trait-based framework to mosses or making comparisons between mosses and vascular plants under any theoretical framework would only be meaningful to the extent that growth form (including lack of roots) and dispersal strategies are considered.

Our findings that trait responses and variability depends on the prevailing environment highlights the limitation of investigating or drawing conclusions about local adaptation from responses to a single environment. Additionally, because phenotypic differentiation may not necessarily have a genetic basis, it is possible in a common garden experiment to confuse or conflate adaptive differentiation arising from phenotypic plasticity with that arising from local adaptation (Gienapp *et al*., 2008).

Finally, there is an on-going taxonomic revision to *S. magellanicum*. The species is considered a complex, comprising at least three species—*S. divinum* and *S. medium* in eastern North America, and *S. magellanicum* sensu stricto in South America (Hassel *et al*., 2018). These species have distinct morphological, molecular and distributional characters. The preliminary study suggests that *S. medium* has an amphi-Atlantic distribution while *S. divinum* is circumpolar in its distribution. Since the pigmentation of “*S. magellanicum”* (as we currently know it) lacks genetic basis (Yousefi *et al*., 2017) and considering the pattern of distribution of these species relative to our field site in Southern Ontario, it is unlikely that we sampled across a mix of *S. medium* and *S. divinum* in a way that would bias our findings. Also, considering that origin had little effect on trait variability, a more likely scenario is that we sampled one species or the other. However, because further study is required on the distribution and identification of these subspecies (Hassel *et al*., 2018), we are unable to accordingly characterize our species and therefore maintain the name *S. magellanicum* for the purpose of this study.

## Conclusion

In summary, we explored the magnitude and pattern of trait variability in *S. magellanicum* from contrasting habitats in the context of phenotypic plasticity and local adaptation. We found that the trait responses were due largely to phenotypical plasticity with little influence on whether plants originated from hummocks or hollows. We also found that trait variability depends on the prevailing light or moisture environment. However, most trait variation remained unexplained by our experimental treatments. Collectively, our results suggest that using traits to draw inferences about the ecology of *Sphagnum* would require an understanding of the mechanisms driving traits and the pattern of trait variability. Lastly, because morphological integration may have an overriding influence on growth traits, we are unclear about the circumstances under which local adaptation might occur or benefit this plant group. We hope that future studies will further explore this area of inquiry in mosses, with consideration for their growth form and recruitment strategies.

## Acknowledgments

The authors thank Mike Mucci and Tannis Slimmon for their technical and material supports in the phytotron. We also thank Sarah McDonald and Rebecca Evans for their assistance with greenhouse and lab measurements. This work was supported by a NSERC Discovery grant to MRT.

## Author contributions

TAO conceived the idea, designed the experiments, collected and analyzed the data. TAO, MRT, DJW and AJS contributed to data interpretation and writing of the manuscript.

